# Horizontally transferred glycoside hydrolase 26 may aid hemipteran insects in plant tissue digestion

**DOI:** 10.1101/2024.04.15.589584

**Authors:** Hunter K. Walt, Seung-Joon Ahn, Federico Hoffmann

## Abstract

Glycoside hydrolases are enzymes that break down complex carbohydrates into simple sugars by catalyzing the hydrolysis of glycosidic bonds. There have been multiple instances of adaptive horizontal gene transfer of genes belonging to various glycoside hydrolase families from microbes to insects, as glycoside hydrolases can metabolize constituents of the carbohydrate-rich plant cell wall. In this study, we characterize the horizontal transfer of a gene from the glycoside hydrolase family 26 (GH26) from bacteria to insects of the order Hemiptera. Our phylogenies trace the horizontal gene transfer to the common ancestor of the superfamilies Pentatomoidea and Lygaeoidea, which include stink bugs and seed bugs. After horizontal transfer, the gene was assimilated into the insect genome as indicated by the gain of an intron, and a eukaryotic signal peptide. Subsequently, the gene has undergone independent losses and expansions in copy number in multiple lineages, suggesting an adaptive role of GH26s in some insects. Finally, we measured tissue-level gene expression of multiple stink bugs and the large milkweed bug using publicly available RNA-seq datasets. We found that the GH26 genes are highly expressed in tissues associated with plant digestion, especially in the principal salivary glands of the stink bugs. Our results are consistent with the hypothesis that this horizontally transferred GH26 was co-opted by the insect to aid in plant tissue digestion and that this HGT event was likely adaptive.

## 1. Introduction

Glycoside hydrolases are a large class of enzymes that break down complex carbohydrate molecules by hydrolyzing glycosidic bonds. There are over 150 known families of glycoside hydrolases, and there is a wide range of substrate specificity across the families (Garron and Henrissat, 2019; Henrissat, 1991). Glycoside hydrolases are important metabolizers in plants, as complex carbohydrates make up a substantial portion of the plant cell wall, so many glycoside hydrolases are necessary for normal plant growth and elongation (Kfoury et al., 2024; Nazipova et al., 2022). Alternatively, glycoside hydrolases encoded in the genomes of other organisms can be used to break down plant cell walls and access nutrients inside plant cells (He et al., 2022; Rafiei et al., 2021).

Horizontal gene transfer (HGT) is the movement of genetic material between genomes without reproduction (Keeling & Palmer, 2008). Initially, HGT was thought to only happen between prokaryotes, and functional HGT from prokaryotes to eukaryotes was met with skepticism as foreign DNA would have to overcome the fundamental differences in the transcriptional machinery between prokaryotes and eukaryotes, and make it to the nucleus of a germline cell for succesful propagation to the next generation (Keeling & Palmer, 2008). Advances in DNA sequencing technologies, which have allowed for easier detection of HGTs, have helped show that bacterial genes have been domesticated by eukaryotic genomes multiple times, as indicated by the gain of introns, polyadenylation sites, eukaryotic signal peptides, and changing codon bias after their insertion in the new host genome (Da Lage et al., 2013; Wybouw et al., 2016). HGT from microbes to insects has happened multiple times (Li et al., 2022; Nakabachi, 2015) and these HGT events have contributed novel genes in several insect orders.

In turn, these HGT-derived genes have been associated with changes in behavior, increases in metabolic breadth, and allowed insects to feed on plants more effectively (Acuña et al., 2012; Ahn et al., 2014; Dai et al., 2021; Huang et al., 2024; Kirsch et al., 2014; Li et al., 2022; Luan et al., 2015; Pauchet & Heckel, 2013; Shelomi et al., 2016; Wybouw et al., 2016). Many of these HGT events are considered adaptive, as functional studies have shown that some of the genes acquired by HGT increase the fitness of insects (Acuña et al., 2012; Dai et al., 2021; Huang et al., 2024; Kirsch et al., 2022). Along with this, many horizontally transferred genes in insects have undergone expansions in copy number. Interestingly, it has been found that HGT genes in bacteria are more likely to duplicate than genes that were inherited vertically (Hooper and Berg, 2003), and studies of HGT from bacteria to insects show a similar pattern (Li et al., 2011). After HGT, gene duplication can lead to functional diversification through neofunctionalization or subfunctionalization of gene copies, allowing for more evolutionary novelty (Dai et al., 2021; Zhang, 2003). Conversely, some of these genes can be lost through pseudogenization or accumulation of deleterious mutations which can be attributed to random processes or relaxed selection at a particular locus (Lynch & Conery, 2000; Zhang, 2003).

There have been many instances of glycoside hydrolase genes being incorporated into the genomes of insects, fungi, and plants via HGT (Acuña et al., 2012; Da Lage et al., 2013; Kfoury et al., 2024; Kirsch et al., 2014; Pauchet and Heckel, 2013; Shelomi et al., 2014; Shen et al., 2003; Shin et al., 2023; Wheeler et al., 2013; Wybouw et al., 2016). In this study, we characterize an HGT event of a gene encoding for a glycoside hydrolase family 26 (GH26) protein, which are enzymes that break down carbohydrates present in plant cell walls, specifically mannans and galactomannans (Braithwaite et al., 1995; Gao et al., 2023; Patel et al., 2016; Zhang et al., 2014). Galactomannans are highly abundant in the endosperm of plant seeds, particularly in legumes, and functional characterizations of GH26 proteins in bacteria have shown that they efficiently hydrolyze locust bean gum and guar gum, both of which are galactomannan polysaccharides extracted from plant seeds (Bågenholm et al., 2019; Gao et al., 2023; Liu et al., 2020; Malgas et al., 2015; Patel et al., 2016; Sharma et al., 2022; von Freiesleben et al., 2019). This HGT can be traced back to an ancestor of the Pentaomoidea and the Lygaeoidea, two subfamilies within the order Hemiptera that include agriculturally relevant crop pests such as the stink bugs. This HGT was identified in a previous genome study of the large milkweed bug, *Oncopeltus fasciatus*, but genomic resources for an in-depth investigation were lacking at the time (Panfilio et al., 2019). The goal of this study is to investigate this HGT comprehensively. Specifically, we 1-used homology searches to detect GH26 orthologs in other insect taxa, 2-used phylogenetic analyses and tree reconciliation to reconstruct patterns of gene gain and loss, and 3-measured gene expression of GH26s across multiple tissues in five taxa from the Hemiptera. Our results uncovered a complex history of expansion and contraction in the insect GH26 gene family and suggest that this HGT is involved in plant tissue digestion during feeding.

## 2. Methods

### 2.1 Genomic data acquisition and candidate GH26 identification

Genome data packages for the brown marmorated stink bug, *Halyomorpha halys* (GCF_000696795.2), the southern green stink bug, *Nezara viridula* (GCA_928085145.1), the neotropical brown stink bug, *Euschistus heros* (GCA_003667255.2) (Singh et al., 2023), and the redbanded stink bug, *Piezodorus guildinii* (GCA_023052935.1) (Saha et al., 2022) were downloaded using the NCBI datasets command line tool (https://www.ncbi.nlm.nih.gov/datasets/docs/v2/reference-docs/command-line/datasets/). We used the redbanded stink bug annotation that was generated in our previous study (Walt et al., 2023), and the neotropical brown stink bug annotation from Singh et al. (2023). The large milkweed bug (*O. fasciatus*) genome data and annotation (version 1.2) (Panfilio et al., 2019) were downloaded from the USDA i5k workspace website (https://i5k.nal.usda.gov/bio_data/836678). The predicted proteome of each genome dataset was functionally annotated using eggNOG mapper v.2.1.9 in DIAMOND mode (Cantalapiedra et al., 2021), leading to the identification of the GH26 sequences. As a first step to identify putative GH26 orthologs, we used OMA standalone v2.5.0 (Altenhoff et al., 2019). The protein sequences placed in the resulting GH26 orthogroup were used to seed BLASTp searches against the brown marmorated stink bug, southern green stink bug, neotropical brown stink bug, redbanded stink bug, and large milkweed bug proteomes to identify diverged homologs that may have been missed during orthology assignment or functional annotation. Other candidate GH26 sequences were identified in insects and bacteria using BLASTp against the NCBI’s nonredundant (nr) protein database, and tBLASTn against NCBI’s whole genome shotgun (wgs) and transcriptome shotgun assembly (TSA). To determine the approximate number of GH26 genes of an organism that only had data present in the TSA database, we used a single candidate GH26 sequence from its transcriptome and aligned it back to the TSA database of the organism it derived from using tBLASTx. We also downloaded all UniProt-reviewed proteins containing the InterPro glycoside hydrolase family 26 domain (IPR022790), which included fungal sequences. To minimize the impact of low-quality sequences and assemblies, we only used sequences from annotated insect genomes and complete bacterial and fungal proteins for all further analyses.

### 2.2 Phylogenetic Analysis

For our phylogenetic analysis, we removed all duplicate proteins. We aligned the resulting amino acid sequences using the MAFFT v7.490 E-ins-I, G-ins-I, and L-ins-I algorithms (Katoh and Standley, 2013), along with the Muscle v3.8.1551 algorithm (Edgar, 2004). The quality of the resulting alignments was compared using MUMSA v.1.0 (Lassmann and Sonnhammer, 2006), and the alignment with the highest score from MUMSA was used to build the phylogeny. The phylogeny was reconstructed using IQTREE2 v.2.0.7 (Minh et al., 2020; Nguyen et al., 2015), incorporating the ModelFinder tool (Kalyaanamoorthy et al., 2017) to select the best-fitting amino acid substitution model identified by the Bayesian Information Criterion. Branch support was assessed using ultrafast bootstrap, Bayesian inference using abayes, and the SH-alrt algorithms incorporated in IQTREE2 (Anisimova et al., 2011; Hoang et al., 2018; Minh et al., 2013; Shimodaira, 2002). For visual clarity, some redundant and highly diverged bacterial and fungal sequences were removed from the less relevant clade in the phylogeny (i.e. the fungal/bacterial clade), and the analysis was run again. Synteny analysis was conducted manually with the aid of NCBI’s genome data viewer (when available) and each organism’s annotation files. BLAST searches were conducted to identify orthologous flanking genes.

### 2.3 Tissue-Specific Expression

To quantify tissue-specific expression levels of HGT GH26 genes, we downloaded 131 SRA datasets from the brown marmorated stink bug, the southern green stink bug, the redbanded stink bug, the neotropical brown stink bug, and the large milkweed bug where specific tissue dissections were reported. The SRA accession numbers of the datasets we used are listed in **Supplementary Data 1**. For each dataset, we trimmed low-quality reads using Trimmomatic v0.39 (unless the dataset was reported as already trimmed) (Bolger et al., 2014), and measured transcript abundance using Kallisto v0.46.2 (Bray et al., 2016). If a reference transcriptome was not available, transcriptomes were generated from annotation and genome files using GffRead (Pertea and Pertea, 2020) and used as indices for Kallisto. Data visualization was conducted in R (R Core Team, 2020).

## 3 Results & Discussion

### 3.1 Characterization of HGT event

Insect glycoside hydrolases are of interest because of their role in digesting plant tissue. In this study, we used homology searches to trace the origin and evolution of the glycoside hydrolase 26 gene family (GH26) in a sample of insect genomes representative of the order Hemiptera. A previous study reported the presence of GH26 homologs in the brown marmorated stink bug (*H. halys*) and the large milkweed bug (*O. fasciatus*) and inferred that these genes were of bacterial origin (Panfilio et al., 2019). We expanded searches to additional stink bug genomes, such as the southern green stink bug (*N. viridula*), the redbanded stink bug (*P. guildinii*), and the neotropical brown stink bug (*E. heros*) and identified multiple new GH26 bacterial-type genes. We also repeated our search strategy in the brown marmorated stink bug and the large milkweed bug genomes and found additional genes in the large milkweed bug genome relative to the original study. Outside these species, the closest homology matches came from bacteria. Within these genes, we found intron regions and eukaryotic signal peptides, an indication that these genes are eukaryotic instead of bacterial contamination in the genome assemblies (**Figure 1A & B**). The general structure of these genes in all insect genomes is a short coding exon (~50 bp) followed by a short intron (~80-250 bp) and a long coding exon (~1000 bp) (**Figure 1B**). Deviations were observed in this general pattern, but this could be the result of errors in assembly and annotation.

**Figure 1:**
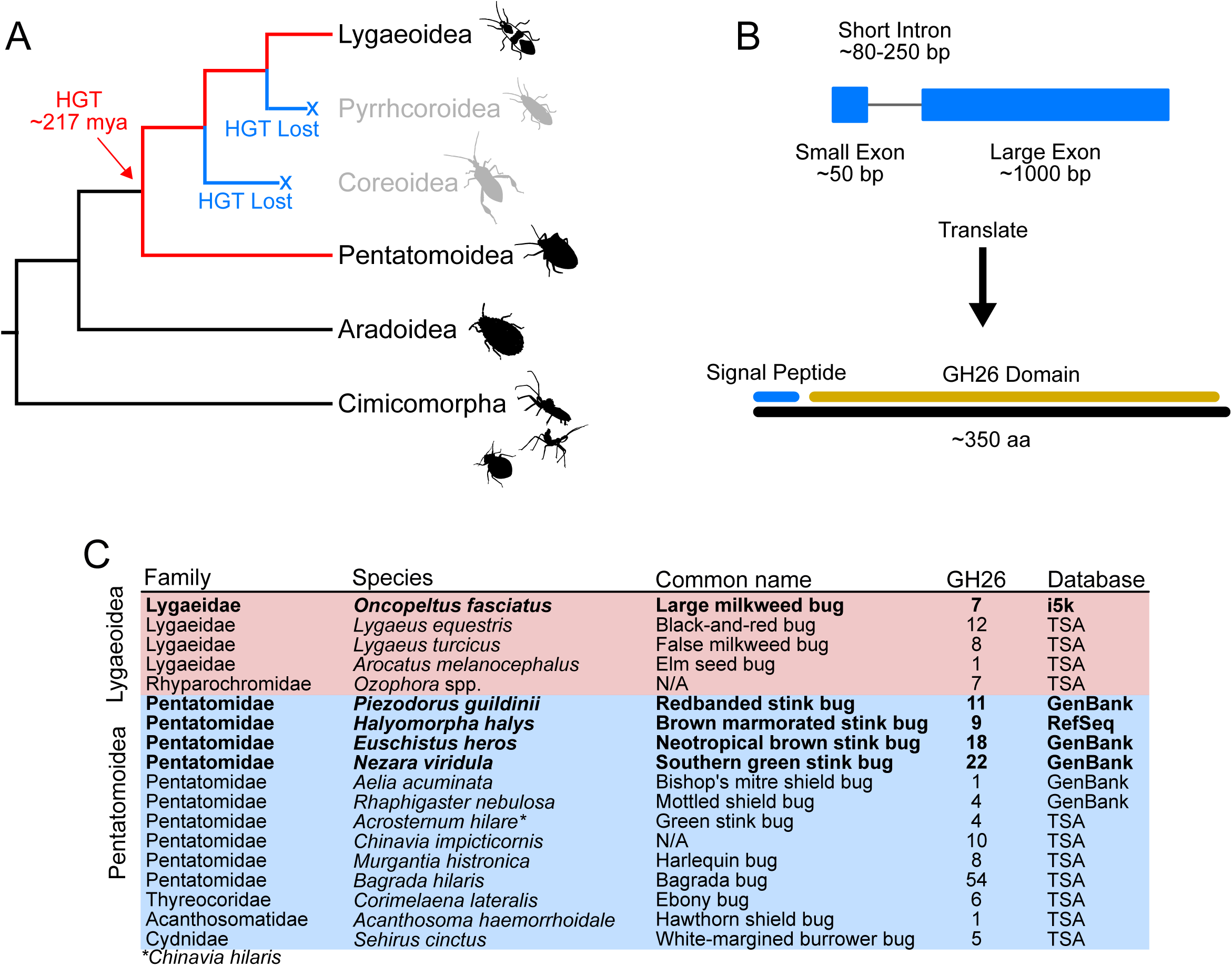
A GH26 gene was horizontally transferred to the common ancestor of the superfamilies Pentatomoidea (stink bugs and shield bugs) and Lygaeoidea (seed bugs and allies) and assimilated to eukaryotic transcription. **A.)** The HGT occurred around 217 million years ago (according to Liu et. al (2019)) and was subsequently lost in the Coreoidea and Pyrrhocoroidea. Phylogenetic relationships used for this tree were derived from Liu et. al (2019). **B.)** Gene and protein structure of insect GH26s. **C.)** Table showing the number of GH26 signatures detected in multiple hemipteran taxa, along with the genomic database we found them in. Taxa in bold were used for subsequent analyses.

We then expanded our searches to all insect lineages. We found that GH26 genes are only present in the superfamilies Pentatomoidea (stink bugs, shield bugs, and their allies) and Lygaeoidea (seed bugs and their allies) (**Figure 1C**). The last common ancestor of the Pentatomoidea and Lygaeoidea superfamilies is shared by the superfamilies Pyrrhocoroidea (bordered plant bugs and red bugs) and Coreoidea (leaf-footed bugs, scentless plant bugs, and allies). Thus, the presence of GH26s in a subset of the groups derived from this ancestor could be the result of independent HGTs specific to each lineage where the genes are present, or alternatively, this could be the product of a single HGT followed by independent losses in the common ancestor of the superfamilies where the genes are apparently absent. To explore this, we estimated phylogenetic relationships for the GH26 candidates identified along with a selection of bacterial and fungal GH26 sequences identified from BLAST searches and the InterPro database. Our phylogeny (**Figure 2**) places all these GH26 genes in a monophyletic group sister to a clade of bacterial GH26s. This suggests that the presence of GH26 genes in the superfamilies Pentatomoidea and Lygaeoidea is the result of a single HGT event of a gene of bacterial origin into the genome of their last common ancestor (i.e. the Trichophora), which is dated to 217 million years ago by Liu et al. (2019). We found no GH26 candidate sequences in the superfamilies Pyrrhocoroidea and Coreoidea which suggests that functional copies of GH26 were secondarily lost independently in these two lineages (**Figure 1A**). Panfilio et al. (2019) had previously reported two GH26 genes (referred to as endo-1,4-beta-mannosidase) in the large milkweed bug genome and nine GH26 genes in the brown marmorated stink bug. Our analyses suggest that there was a larger expansion of GH26 in the large milkweed bug than was previously reported (Panfilio et al., 2019), as we detected seven annotated GH26 copies in its genome relative to the two reported in the original study (**Figure 1C**). We found GH26 genes present in multiple copies in nearly every lineage that it is present, suggesting that the HGT GH26 may be adaptive in these insects (**Figure 1C**).

**Figure 2:**
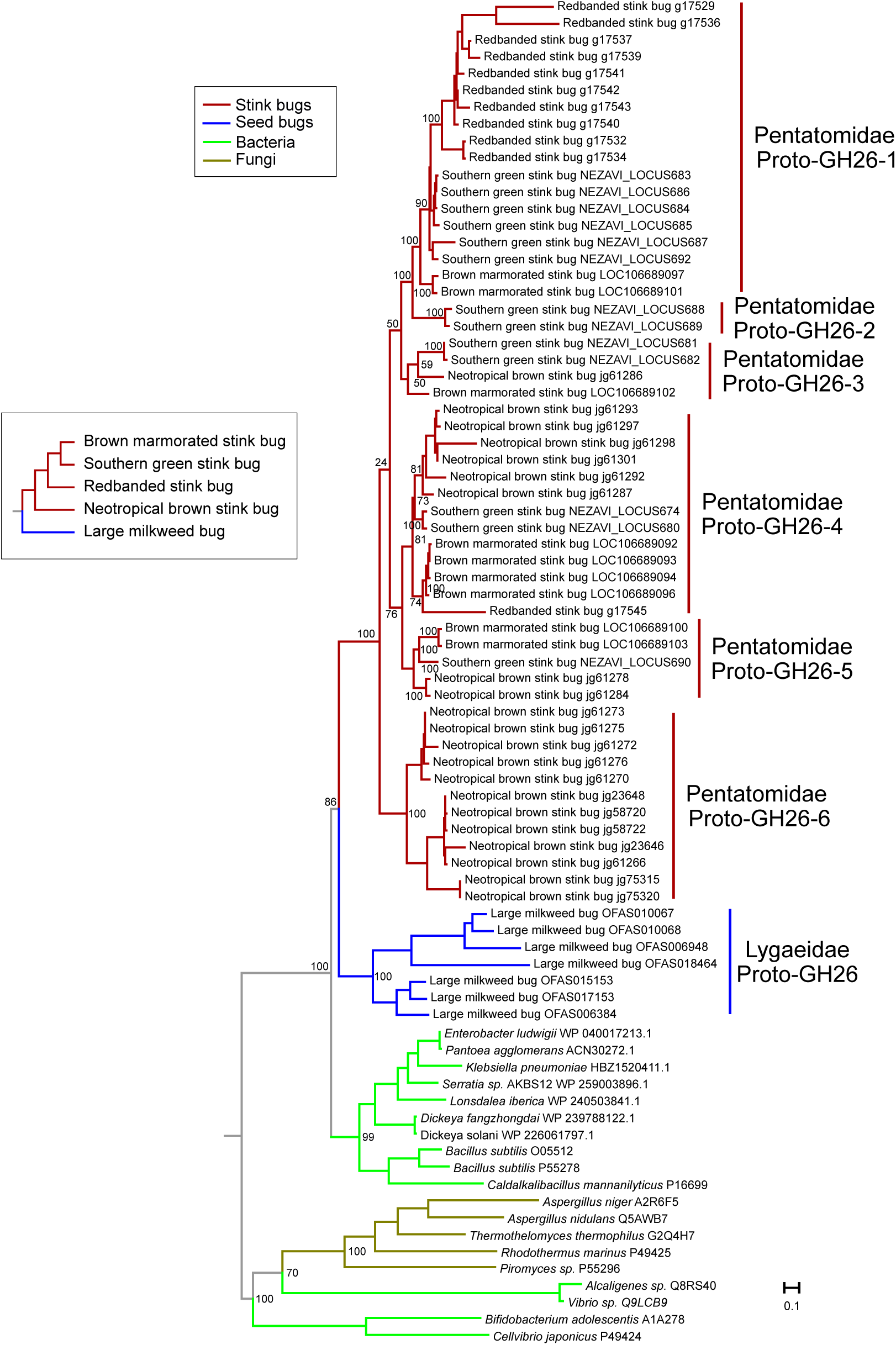
Gene family evolution of GH26 genes in insects. Our phylogeny suggests that a single GH26 gene was horizontally transferred from bacteria and independently expanded in the seed bugs and the stink bugs. A species tree of the insects used in the phylogeny is shown and it is based on a previous phylogenomic analysis that we conducted (Walt et al., 2023). Using the species tree and the GH26 gene tree, we infer at least 6 GH26 gene copies present at the last common ancestor of the Pentatomidae (stink bugs), which are annotated along the tree as “Proto-GH26s”. Because there is only one annotated genome from the family Lygaeidae, we can only infer one proto-GH26 at their common ancestor. UF bootstrap support values are indicated on relevant nodes.

### 3.2 Gene family evolution of the GH26 gene

To investigate lineage-specific patterns of gene gain and loss after the HGT of the GH26 gene, we delved deeper into the phylogeny reported in **Figure 2**. We found that the GH26 sequences cluster by family, with the large milkweed bug sequences placed in a monophyletic group sister to the clade that includes all the stink bug GH26 sequences, suggesting that a single GH26 gene was present in their last common ancestor and expanded independently expanded in these insect families (**Figure 2**). For the large milkweed bug, this gene was duplicated at least 6 times after the HGT event (a total of seven GH26 genes annotated). Because the large milkweed bug is the only annotated genome from the seed bugs (Lygaeidae), we could not infer the number of ancestral genes at this family’s most recent common ancestor. As for the stink bugs (Pentatomidae), our phylogeny suggests that at least 6 GH26 genes were present in their last common ancestor (**Figure 2**) and that the GH26 repertoires of extant lineages have been shaped by a complicated history of duplication and loss (**Figure 2**). This suggests that after the HGT, the GH26 genes followed patterns consistent with the gene birth and death evolution model (Ota and Nei, 1994). Notably, our phylogeny shows another potential HGT of a GH26 gene from bacteria to fungi, consistent with previous findings (Millward-Sadler et al., 1996).

For most lineages, our synteny analysis confirms that these duplications were in tandem, as GH26 genes are generally arranged in clusters flanked by a conserved alpha-(1,6)-fucosyltransferase gene on one side, and genes encoding for a basic helix-loop-helix domain-containing protein and an odorant receptor on the other side (**Figure 3**). This pattern was most clear in the southern green stink bug and the redbanded stink bug, as they have chromosome-level genome assemblies (**Figure 3**). Synteny was difficult to establish in the neotropical brown stink bug and the large milkweed bug, as their genomes are highly fractionated in this area (**Figure 3**). Additional evidence for the birth-and-death pattern of evolution of these genes comes from the presence of multiple pseudogenes in the GH26 clusters.

**Figure 3:**
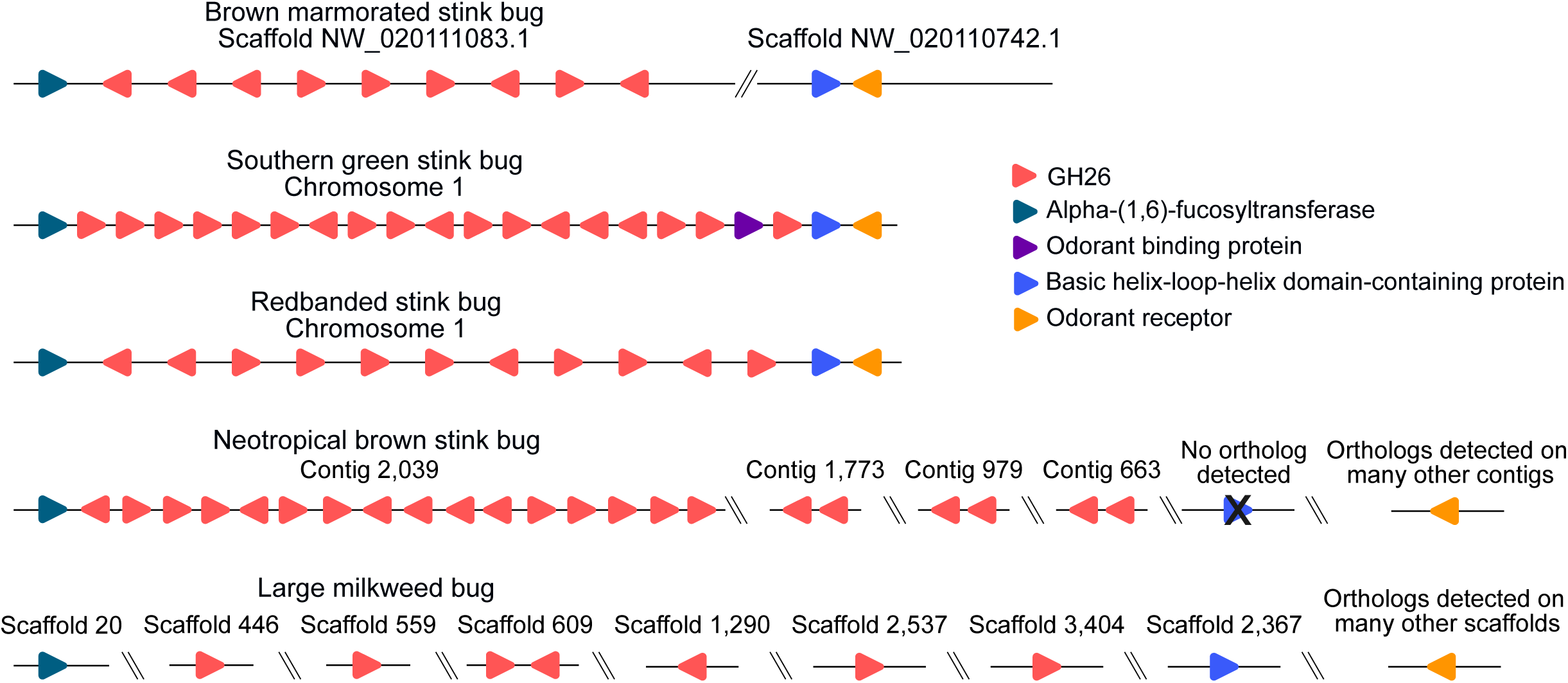
Synteny of GH26 genes across all annotated taxa. Insect GH26 genes form clusters derived from multiple tandem duplications. These clusters are flanked by a gene encoding for an alpha-(1,6)-fucosyltransferase on one side, and a basic helix-loop-helix domain containing protein and an odorant receptor on the other side. Synteny of the neotropical brown stink bug and the large milkweed bug genomes were difficult to determine as they are fragmented in this area.

### 3.3. Expression of GH26 genes in insect tissues

The GH26 family proteins are known to metabolize carbohydrates present in plant cell walls, particularly mannans and galactomannans (Araki et al., 2000; Braithwaite et al., 1995; Gao et al., 2023; Okazaki et al., 2002; Taylor et al., 2005; Zhang et al., 2014), so we hypothesize that the HGT insect GH26s play a role in plant cell wall degradation. Thus, we expected them to be highly expressed in digestive tissues. For phytophagous hemipteran insects, digestion starts extra orally, as they use their piercing-sucking mouthparts to probe plants and excrete potent saliva that begins to digest plant tissues (Backus et al., 2005; Liu and Bonning, 2019; Lomate and Bonning, 2016; Marshall et al., 2023). After this, the digested plant tissues are ingested into their gut where they are further metabolized (Lomate and Bonning, 2016; Miles, 1972). Because of this, we expect the HGT GH26 genes to be differentially expressed in the salivary glands and the gut. For the southern green stink bug, the redbanded stink bug, and the brown marmorated stink bug, we found that expression of all copies of GH26 is salivary biased, with extremely high expression in the principal salivary gland (PSG), where most of the digestive enzymes are expressed (Liu and Bonning, 2019) (**Figure 4**). Many stink bugs are agricultural pests, as they feed on the seeds of soybean and other important crops (Corrêa-Ferreira and De Azevedo, 2002; Depieri and Panizzi, 2011). Interestingly, the endosperm of the seeds of legumes is rich in galactomannans, a substrate of many GH26 proteins (Sharma et al., 2022). This suggests that the HGT GH26s derived from the PSG of stink bugs may be involved in extra-oral digestion of seed tissues and could be of interest to agriculture, although functional validation is necessary.

**Figure 4:**
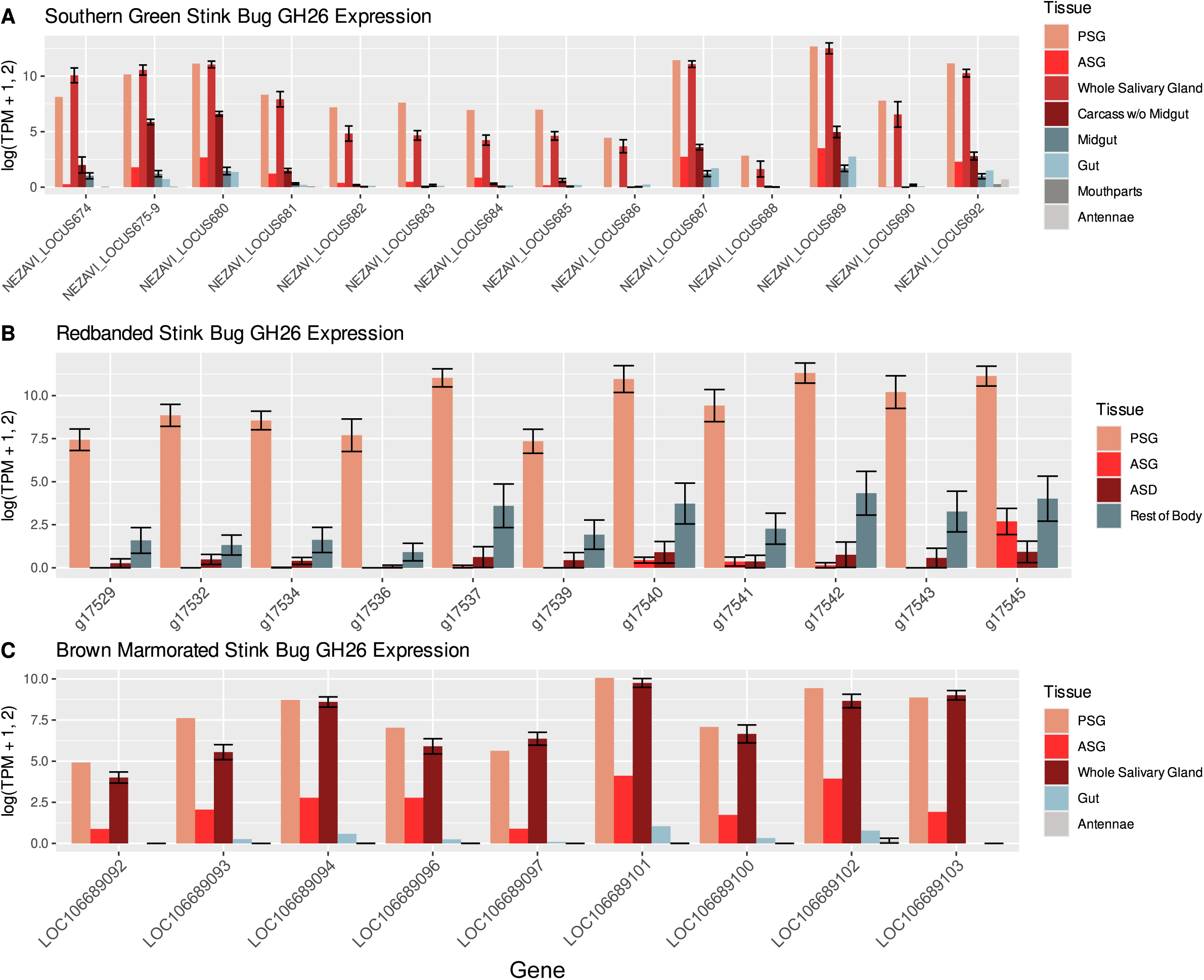
Expression of GH26s in the stink bugs. All samples containing salivary tissues are in shades of red, and all samples containing gut tissues are in shades of blue. All other tissues are in shades of grey. Error bars represent the standard error of the mean (SEM) and are shown when multiple replicates are present within a tissue. **A.)** Expression of southern green stink bug GH26 genes across salivary, gut, mouthparts, and antennae tissues. Loci 675-679 are shown as one locus because their sequences are identical. **B.)** Expression of redbanded stink bug GH26 genes across salivary tissues, and the rest of the body without salivary tissues. **C.)** Expression of brown marmorated stink bug GH26 genes across salivary, gut, and antennal tissues. The highest expression values are consistently in the principal salivary gland. TPM = Transcripts per million, PSG = principal salivary gland, ASG = accessory salivary gland, ASD = Accessory salivary duct.

Unfortunately, there are no RNA-seq datasets of salivary tissues for the neotropical brown stink bug or the large milkweed bug, thus we included these analyses as supplemental (**Supplementary Figures 1 & 2**). Interestingly, the observed patterns are consistent with our predictions. In the neotropical brown stink bug, the GH26 genes are expressed in the gut tissues and the fat body, another site of carbohydrate metabolism (Arrese and Soulages, 2010) (**Supplementary Figure 1**). In the large milkweed bug, we found high expression of GH26 genes in the gut, head, and thorax (**Supplementary Figure 2**). The head and thorax of the large milkweed bug could feasibly have contained salivary tissues, although we cannot confirm this. Interestingly, three GH26 genes in the large milkweed bug (OFAS010067, OFAS010068, and OFAS018464) have biased expression in the gut, a pattern that we never observed in the stink bugs (**Supplementary Figure 2, Figure 4**). Furthermore, the large milkweed bug gene OFAS006948 exhibited no expression at all, suggesting that it might have become a pseudogene (**Supplementary Figure 2**). Overall, there was low expression of these GH26 genes in non-digestive/metabolic tissues (**Figure 4, Supplementary Figures 1 & 2**). There have been many other instances of genes from other glycoside hydrolase families being horizontally transferred from bacterial and fungal genomes to insects, which suggests that HGT GH26 genes could aid plant tissue digestion in the Pentatomoidea and the Lygaedoidea (Acuña et al., 2012; Kirsch et al., 2014; Pauchet and Heckel, 2013; Shelomi et al., 2014; Shen et al., 2003; Shin et al., 2023; Wheeler et al., 2013; Wybouw et al., 2016).

## 4. Conclusions

Our work investigates the HGT of a GH26 gene from bacteria to the common ancestor of the stink bugs and the seed bugs around 217 million years ago (Liu et al., 2019). After the HGT, the GH26 gene was domesticated by the insect genome by gaining introns and eukaryotic signal peptides. Then, the GH26 gene followed complex patterns of duplication and loss consistent with birth-and-death evolution and is only present in the superfamilies Lygaeoidea and Pentatomoidea (**Figure 1**, **Figure 2**). In extant taxa, the GH26 gene mostly exists in high copy numbers, and we inferred that there were at least six copies of the GH26 gene in the last common ancestor of the stink bugs (**Figure 1C**, **Figure 2**). Furthermore, the HGT GH26 gene exhibits biased expression to digestive tissues (**Figure 4, Supplementary Figures 1 & 2**), especially in the principal salivary glands of stink bugs (**Figure 4**), some of which are significant crop pests. Thus, we propose that after HGT, GH26 was co-opted by insects to feed on their plant hosts more efficiently. Specifically, we predict that insect GH26s are important for digesting seed tissues because bacterial GH26s efficiently hydrolyze galactomannan-rich substances such as the endosperm of seeds (Bågenholm et al., 2019; Gao et al., 2023; Liu et al., 2020; Malgas et al., 2015; Patel et al., 2016; Sharma et al., 2022; von Freiesleben et al., 2019). This is consistent with our results as many of the taxa that retained GH26s feed on the seeds of plants, such as the seed bugs and many stink bugs (Corrêa-Ferreira and De Azevedo, 2002; Depieri and Panizzi, 2011; Feir and Beck, 1963). Functional studies of insect GH26s would be useful to assess their true role in insect feeding and fitness.

## Acknowledgments

The authors would like to thank Chris Bass and Benjamin Hunt for providing us with their annotation of the neotropical brown stink bug genome.

## Author Contributions

**Hunter K. Walt:** Conceptualization, Methodology, Software, Validation, Formal Analysis, Investigation, Data Curation, Visualization, Writing - Original Draft, Writing - Review & Editing. **Seung-Joon Ahn:** Conceptualization, Methodology, Validation, Investigation, Writing - Review & Editing. **Federico G. Hoffmann:** Conceptualization, Methodology, Supervision, Writing - Review & Editing

