## Supplementary Figures 1 & 2 for "Horizontally transferred glycoside hydrolase 26 may aid hemipteran insects in plant tissue digestion"

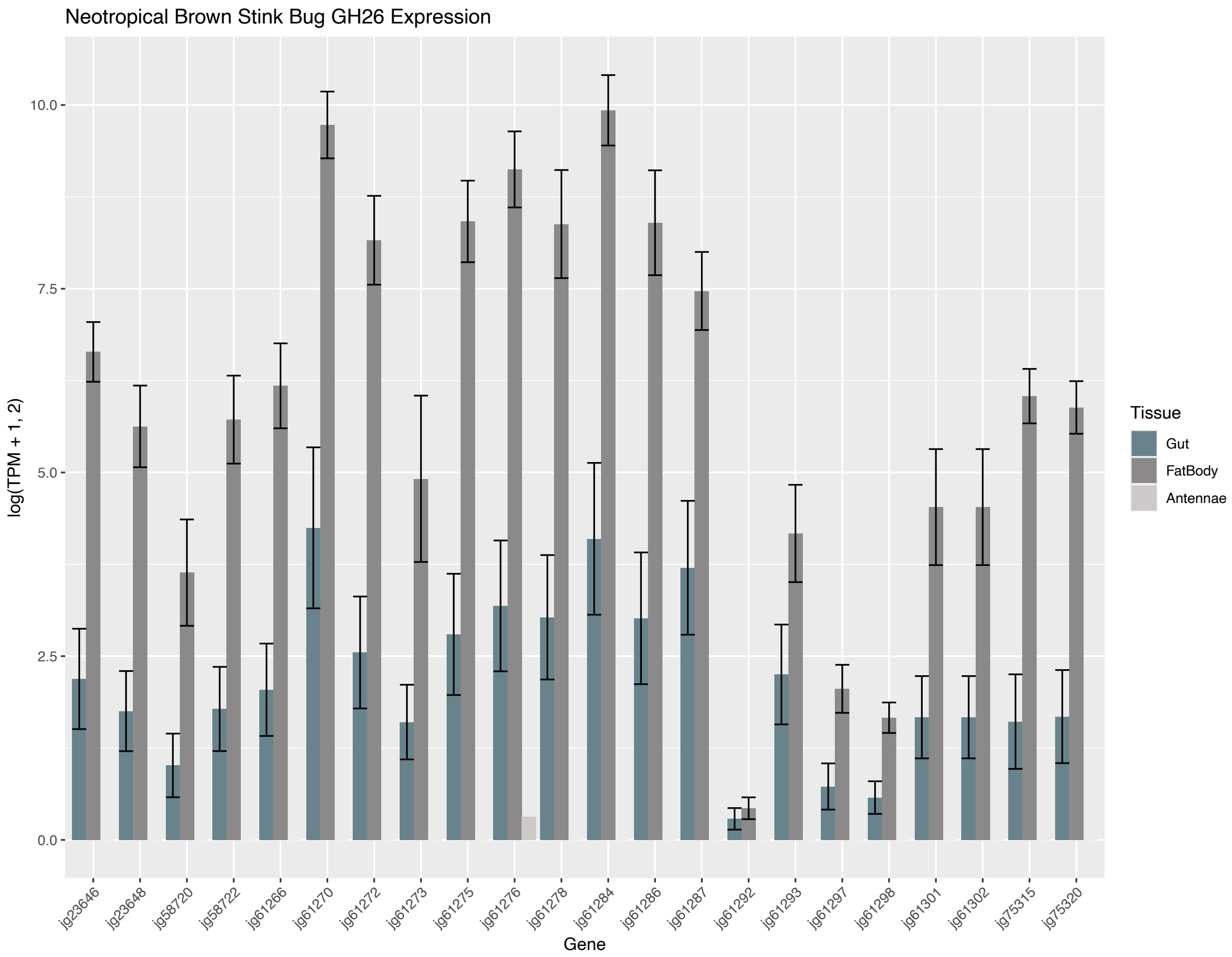


**Supplementary Figure 1:** Expression of GH26 genes in the gut, fat body, and antennae of the neotropical brown stink bug, *Euschistus heros*.

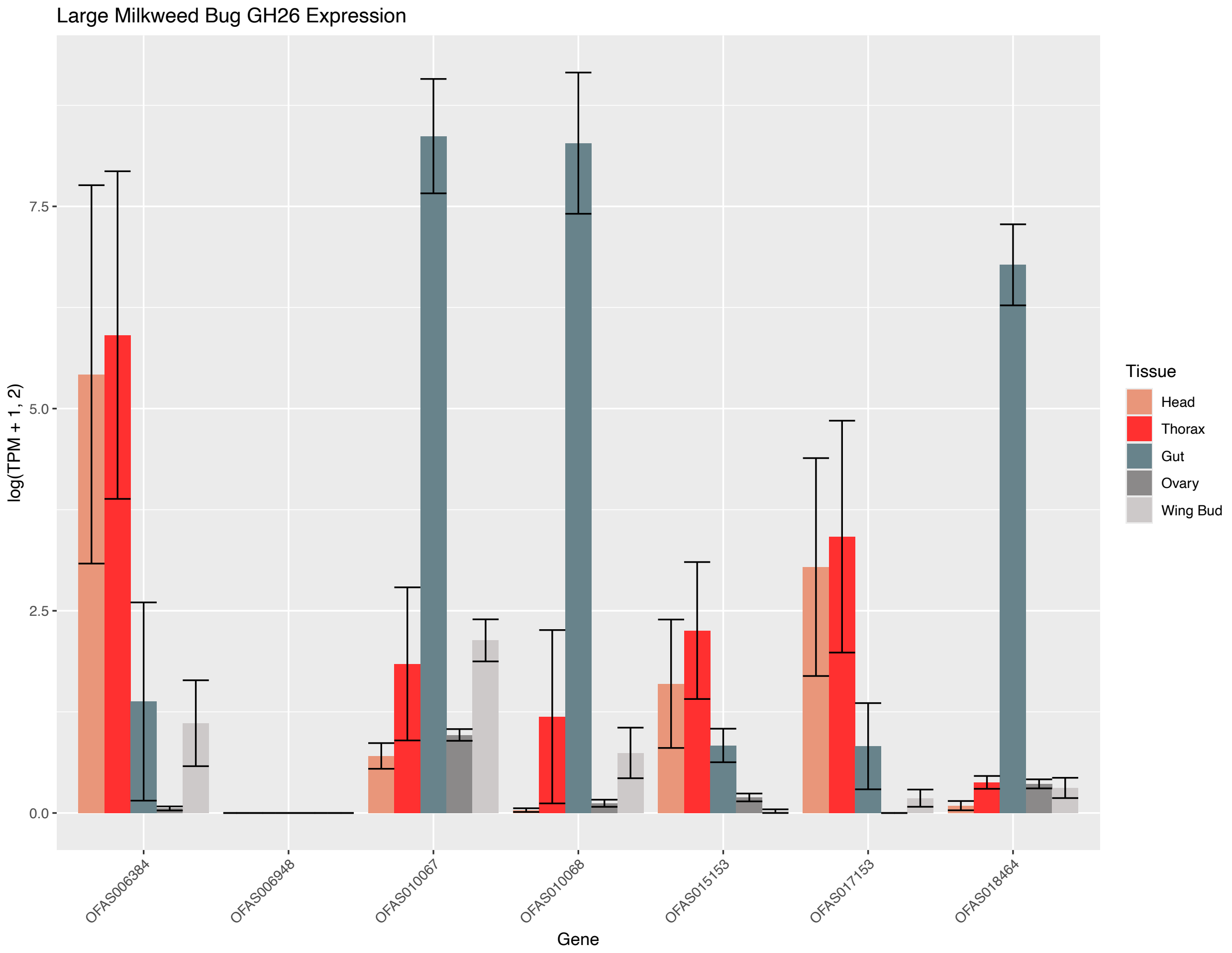


**Supplementary Figure 2:** Expression of GH26 genes in the head, thorax, gut, ovary, and wing bug of the large milkweed bug, *Oncopeltus fasciatus*
